# SAbDab2: The structural antibody database in the age of machine learning

**DOI:** 10.64898/2026.06.16.732554

**Authors:** Henriette L. Capel, Odysseas Vavourakis, Benjamin H. Williams, Christopher R. Taylor, Charlotte M. Deane

## Abstract

The Structural Antibody Database (SAbDab) is a publicly available repository of experimentally determined antibody structures, first released in 2013. Explicit support for single-domain antibodies was added in 2021, with SAbDab-nano. Recently, increasing interest in antibodies has led to a proliferation of novel antibody formats, while simultaneous advances in machine learning have increased demand for standardised, high-quality structure data. Here, we present SAbDab2, re-engineered for the machine-learning age. It introduces support for a variety of new formats, and makes it easy to retrieve and compare all known structures of a given antibody. In addition, SAbDab2 provides ready access to ML-grade structures of antibody and antibody–antigen-complexes, with standardised, versioned train/test splits. These will be updated every six months going forward, and are available at https://zenodo.org/records/20083995. SAbDab2 itself is updated weekly and is freely available at https://sabdab2.opig.stats.ox.ac.uk.

## Introduction

Since its release in 2013, the Structural Antibody Database (SAbDab) (1) has served as the primary reference for antibody structure data. Consistently maintained and updated weekly since its inception, it aims to be a comprehensive repository of all experimentally determined antibody structures in the public domain, annotated on sequence, structure, and functional levels. Explicit support for single-domain antibodies (SAbDab-nano) was added in 2021 (2). Detailed annotations and consistent maintenance have made SAbDab a central resource supporting important advances in the field. For example, SAbDab has been used to study antibody-antigen interactions (3–5), including SARS-CoV-2 (6, 7); to predict antibody structure (8–10); to design antibodies *de-novo* (11–14); and to investigate antibody flexibility (15, 16).

In recent years, experimental advances in solving protein structures (17) and the growing success of antibodies as therapeutics (18) have expanded the known antibody structural space^1^. These developments have also driven growing interest in alternative antibody formats and constructs, such as multi-specific antibodies and antibody fragments (19, 20), expanding their applications and highlighting improvements in developability and manufacturability (20). Of the 26 antibodies in regulatory review at the end of 2025, 38% had a non-conventional format (18). The need for systematic and comprehensive annotation of such constructs is increasingly pressing.

In parallel, emerging computational technologies have led to substantial advances in protein structure and complex prediction (21), with a variety of deep learning approaches now available for antibody structure prediction, antibody–antigen complex modelling, and *de novo* antibody design (9, 14, 22). The success of such models hinges on high-quality data, carefully partitioned into train and test sets to avoid data leakage (23). Fair and meaningful model comparison is predicated on these data splits being standardised (24).

Here, we present SAbDab2, a comprehensive restructuring of SAbDab designed to systematically annotate structures of a wide array of antibody formats for use in machine-learning (ML) applications. As part of this overhaul we introduce SAbDab2 IDs, which uniquely identify single- and paired-chain variable regions by their IMGT numbering. These group structures with identical variable-domain sequences together across different PDB IDs, bound states, formats, and constructs (see Figure 1), enabling direct comparison of *apo* and *holo* conformations, facilitating epitope analysis, supporting investigation of antibody flexibility, and simplifying redundancy filtering. We also describe our updated processing pipeline, including antibody identification and numbering with ANARCII (25), support for the PDBx/mmCIF format (26), and refined definitions of antigen pairing. Alongside SAbDab2, we introduce a high-quality, curated subset of structures, post-processed specifically with ML in mind, alongside standardised, versioned train/test-splits for tasks like structure prediction and antibody–antigen complex modelling.

**Fig. 1.**
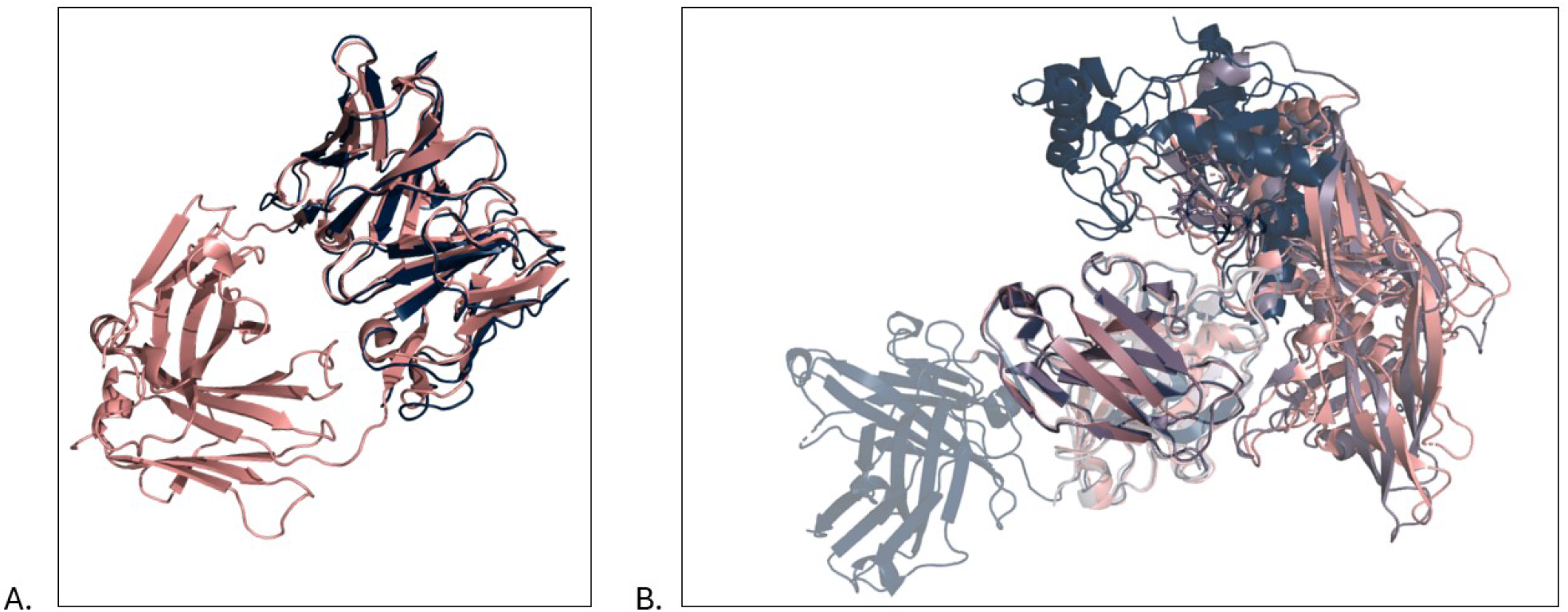
SAbDab2 IDs group together antibody structures with identical variable-domain sequence numbering. (A) A single SAbDab2 ID can attach to antibody structures of different types and constructs if all variable-domain numerable segments are identical. For example, the Fv structure composed of pdb_00006vrl chains “B” and “C” (dark blue) and the Fab structure comprising pdb_00005i66 chains “A” and “B” (pink) are assigned to SAbDab2 ID sabdab2_H0142L00YY. (B) SAbDab2 IDs can reveal different possible antigens, as shown for sabdab2_H0043L003X. Here, a Fab structure (pdb_00003ru8; dark blue) and two Fv structures (pdb_00008sxi; pink, pdb_00009nc3; purple) share the same Fv sequence but bind different antigens. In pdb_00003ru8, (heavy) chain “H” and (light) chain “L” are in close in close proximity to a protein named “Epitope Scaffold 2bodx43” (chain “X”) and a hapten (chain “A”). Their counterpart chains “G” and “H” in pdb_00008sxi bind “Envelope glycoprotein gp160” (chain “E”), while in pdb_00009nc3 chains “M” and “N” are in close proximity to “Envelope glycoprotein gp120” (chain “E”) and a sugar (chain “p”). All antibodies are aligned on their heavy and light framework residues and visualised with PyMol (30).

Unlike other recent antibody structure datasets, such as SAAINT-DB (27), SNAC-DB (28) and NAStructuralDB (29), SAbDab2 is the only tool to provide a web interface; user-directed query and download facilities; and curated, ML-ready data with standardised splits.

SAbDab2 is updated weekly and freely accessible at https://sabdab2.opig.stats.ox.ac.uk. Versioned, ML-ready data splits are publicly available for download at https://zenodo.org/records/20083995, and new versions will be generated at least once every six months.

## Results

SAbDab2 is a redesigned version of the Structural Antibody Database (SAbDab), built to to support and drive the creation of ML models. For more than a decade, SAbDab has served as the primary reference for structural antibody data. This update responds to developments in the field and enables the direct use of this foundational resource in modern computational workflows.

### SAbDAb2 identifies antibody numerable domains using ANARCII

The data processing and annotation pipeline is visualised in Figure 2. For a detailed description see Methods. The publicly available antibody structures in SAbDab2 are scraped from the PDB (31), by downloading the PDBx/mmCIF files of all protein entries. This has been the standard PDB deposition format since July 2019 (26), and is now the only available format for many entries. PDB entries are accessed by their 12 character identifiers (e.g. “pdb_00001abc” instead of “1abc”), anticipating changes to the PDB slated for 2028^2^. Sequences are extracted from FASTA files davailable on the PDB, and numberable sequence segments identified with ANARCII (25), a language model-based antibody numbering tool. Unlike its predecessor AN-ARCI (32), used in SAbDab up to this point, ANARCII enables enables identification of alternative antibody formats such as the shark-derived single-domain VNARs (25).

**Fig. 2.**
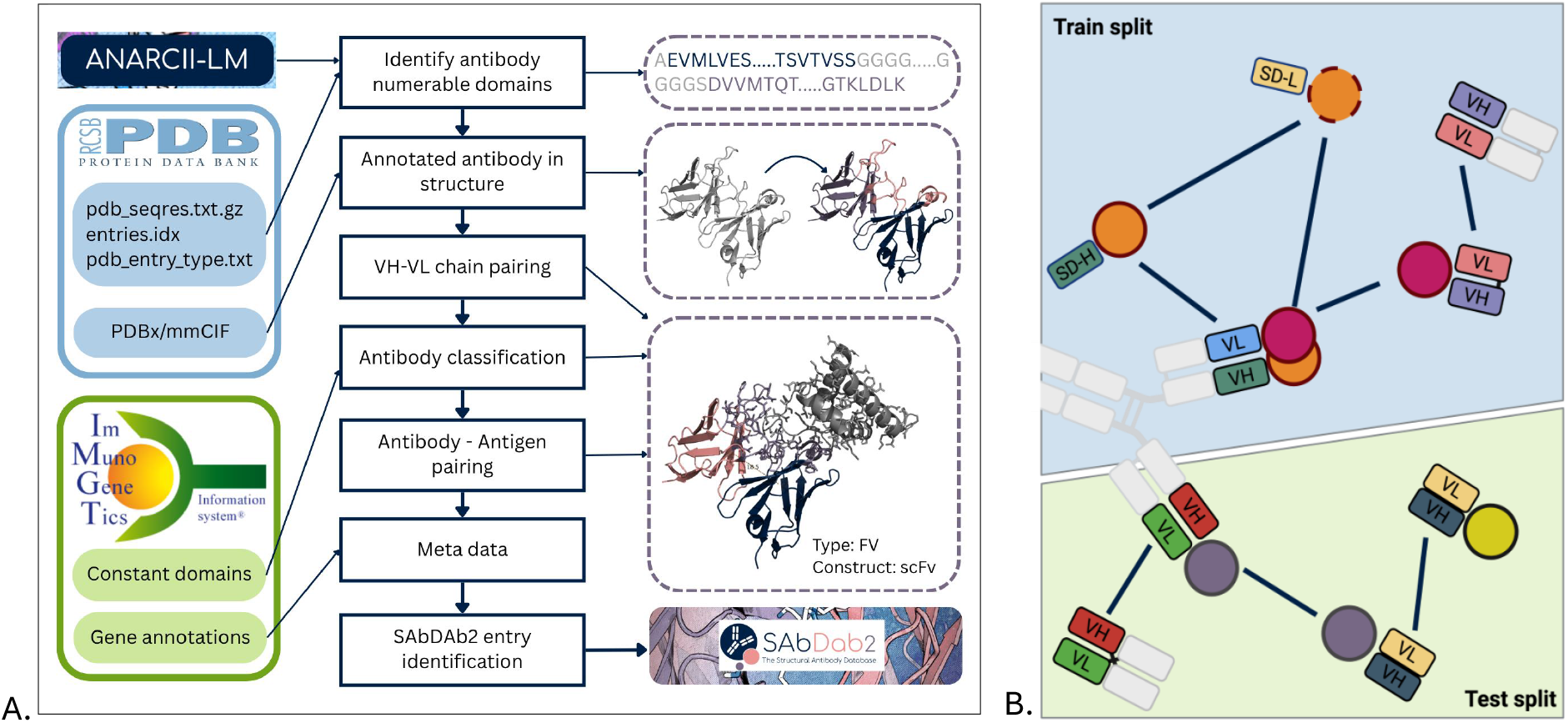
Data processing pipepines for SAbDab2. (A) SAbDab2’s data processing pipeline for the curation, filtering, annotation of structural antibody data. Sequence and strucural data is extracted from the PDB and antibody instances are identified using ANARCII. Author chain IDs are split in the mmCIF files when multiple antibody numerable segments are present in one chain. Identified VH and VL chains are paired based on a close proximity of conserved framework regions, after which antibody types and constructs are determined. Antigens are than classified and paired with the antibody instances based on close proximity. Gene annotations are derived from the IMGT and SAbDab2 IDs are assigned based on the IMGT numbered VH and/or VL sequence. (B) Schematic illustration of the ab-ag-split. High-quality structures of individual antibody instances are split into a standardised train/test split based on sequence similarity. Instance structures are grouped together (connected by edges) if their antibody or any antigen chain fall into a common cluster (indicated by colour). Connected components of the resulting sparse graph are then allocated to the train and test partitions. This ensures similar antibodies and antigens are always grouped together into either train or test, while allowing us to separate distinct antibodies that happen to co-occur in the same PDB entry.

### SAbDab2 IDs enable structure comparisons across PDB IDs

Prior versions of SAbDab organised antibody structure data by PDB accession ID, each of which could contain several copies of multiple sequence-distinct antibodies. SAbDab2 distinguishes antibody instances: individual variable-region structures, comprising either a single domain or a heavy–light pair. These instances are then assigned SAbDab2 IDs, which group together antibody instances with identical variable sequences across PDB IDs (e.g. “sabdab2_H0142L00YY” with heavy-chain numbering “H0142”, and light-chain numbering “L00YY”). These IDs facilitate the study of different structural conformations, antibody formats (see Figure 1A), and antigen targets (see Figure 1B).

Figure 3 shows the cumulative number of antibody instances and PDB IDs represented in SAbDab2, and the corresponding number of unique antibodies (SAbDab2 IDs), by PDB release year. As of 27 May 2026, SAbDab2 contains 21 237 distinct antibody instances (unique structures), derived from 11 085 PDB IDs, and corresponding to 6 540 SAbDab2 IDs (unique variable regions).

**Fig. 3.**
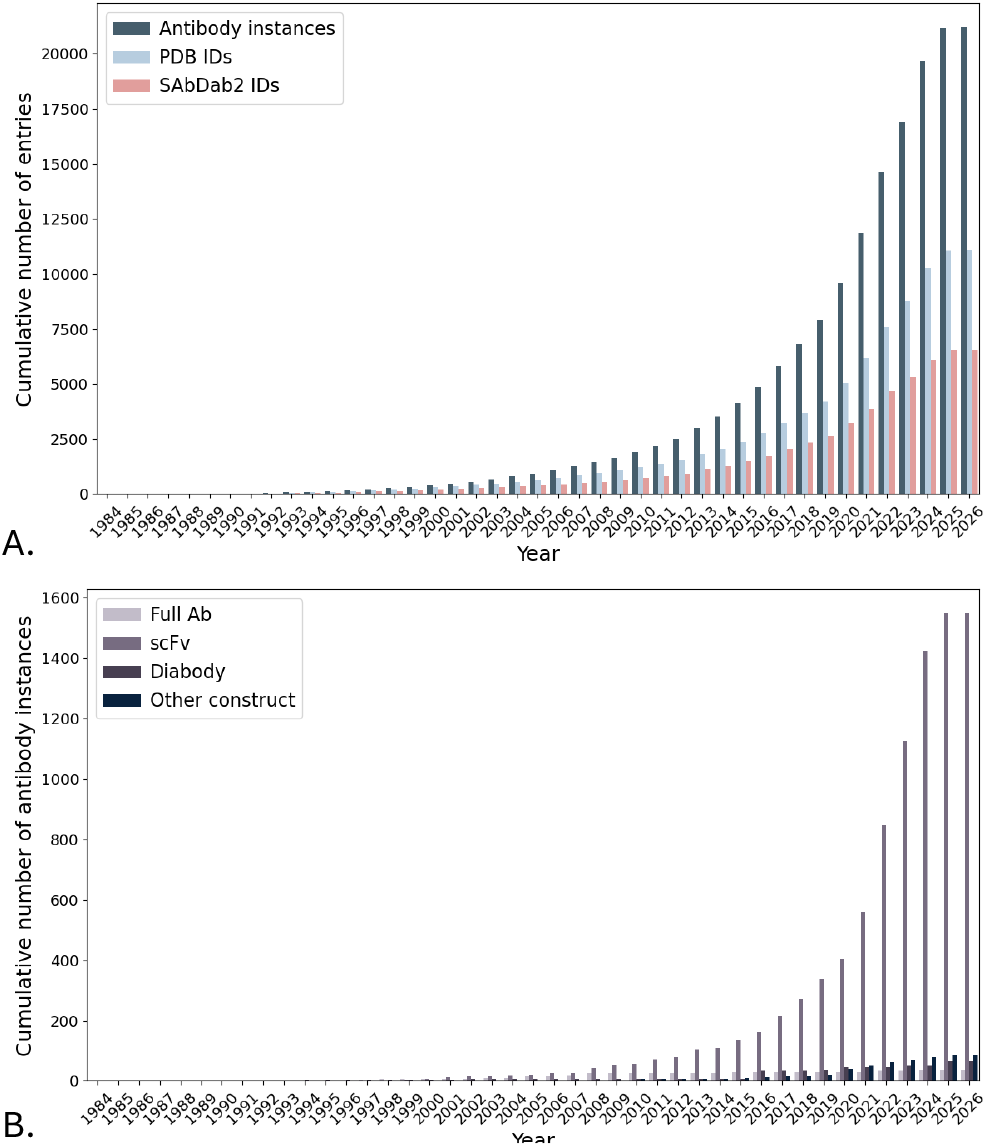
SAbDab2 dataset statistics. (A) The cumulative number of antibody instances and PDB IDs represented in SAbDab2, and the corresponding number of unique antibodies (SAbDab2 IDs), by PDB release year. Numbers for 2026 cover the time until 27 May 2026. The number of instances in the PDB falls behind the amount of structural data available for unique antibody sequences. (B) The cumulative number of antibody instances of the constructs “Full Ab” (light purple), “scFv” (purple), “Diabody” (dark purple), and “Other” (navy) released in the PDB by the end of the year. This shows that engineered constructs are more observed in recent years.

### SAbDab2 annotates engineered antibody formats and constructs

Engineered antibody constructs are increasingly common, frequently combining multiple numerable immunoglobulin domains within a single polypeptide chain (33). Heavy- and light-chain pairing may occur within one physical chain (e.g., scFvs) or across chains (e.g., diabodies). To correctly process these types of entries, SAbDab2’s data processing pipeline systematically splits numberable domains into separate logical chains. For example, domains originally assigned to chain “A” are relabeled as “A1” and “A2” (see Figure 2A). With these changes, all numberable segments in the structure can be identified and numbered. SAbDab2-parsed structure files contain these new chain labels; the original standardised mmCIF chain identifiers (“label_asym_id”), and residue numbering (“label_seq_id”) are preserved.

Numerable sequence segments with resolved structure coordinates are paired based on proximity of the conserved residues at IMGT position 104, forming antibody instances. Instances are then classed into types: “FV”, “FAB”, “FAB+FC”, “SD-H”, “SD-L”, or “VNAR”. Single-domain antibodies (“SD-H”, “SD-L”, and “VNAR”) form a dedicated subset of SAbDab2, SAbDab2-nano, but are also accessible through the main data portal.

Antibody instances of any antibody type can occur within constructs, also with defined types. For example, an “FV” can have (engineered) construct type “SCFV”. Construct types and the antibody instances comprising any given construct are systematically annotated. SAbDab2 currently stores 1 551 scFv and 64 diabody structures, alongside 19 complete antibodies (comprising 38 variable domains; see Figure 3B).

### SAbDAb2 identifies functional antibody-antigen complexes

As in SAbDab, SAbDab2 assigns antigens to antibody instances based on spatial proximity to CDR residues. Author-assigned antigen chains are systematically split by molecular type to enable more detailed complex characterization. To focus on biologically relevant interactions, antigen assignments are restricted—where available—to author-defined biological assemblies. SAbDab2 also newly allows antibodies to be annotated as antigens. Of the total 21 237 antibody instances, SAbDab2 contains 18 907 *holo* structures, including 15 925 involving to at least one protein chain assigned as antigen.

### SAbDab2’s architecture allows for continuous updating and maintenance

SAbDab2 updates weekly, in a semi-automated fashion. The pipeline flags uncertain annotations for manual inspection, and allows for intervention to correct or restore misannotations flagged by the database maintainers or the community.

### SAbDab2 is an easily accessible database of all publicly available antibody structures

SAbDab2 may be accessed at https://sabdab2.opig.stats.ox.ac.uk where the data can be downloaded and searched by PDB ID, SAbDab2 ID, structure detail, experimental detail, sequence similarity, and more (see Figure 4). Search results (except for searches for specific SAbDab2 IDs) are presented by PDB ID by default, but can also be organised by SAbDab2 ID. Both IMGT-numbered antibody structures and IMGT numbered antibody–antigen complex structures are available, in addition to bulk downloads and summary files.

**Fig. 4.**
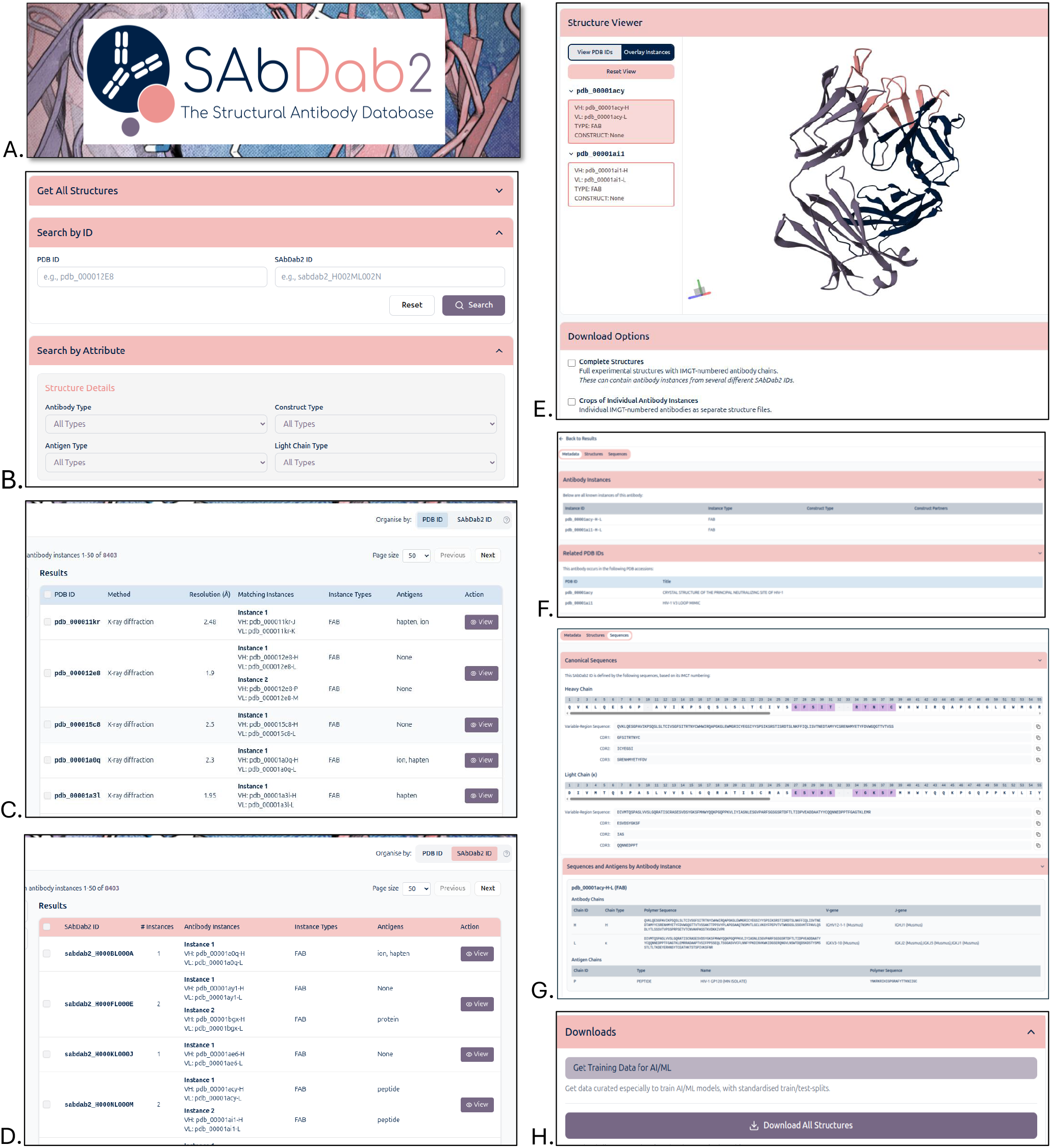
Overview of the SAbDab2 website. (A) The SAbDab2 logo. (B) The SAbDab2 search page allows to search all structures, a specific PDB or SAbDab2 ID, and search by attribute. (C). An example result page organised by PDB ID. (D) An example result page organised by SAbDab2 ID. (E) The structure viewer interface enables visualisation of single antibody instances within the entry or all entries. For the SAbDab2 ID page antibody instances can be aligned on their variable domain framework regions. (F) An example of the “Metadata” tab of a SAbDab ID page. Entry information is shown per antibody instance and the “Related PDB IDs” allows you to navigate to the SAbDab2 PDB ID pages that are part of the SAbDab2 ID. The “Related SAbDab2 IDs” list all SAbDab2 IDs containing the same heavy or light numbered sequence as the viewed SAbDab2 ID. (G) An example of the “Sequence” tab of a SabDab ID page, showing the IMGT numbered sequence and the antibody information per instance. (G) The full database, a filtered database, and specific entries can be downloaded from SAbDab2. A link to the versioned and standardised splits can also be found on the website.

Related structural antibody databases are SAAINT-DB (27), SNAC-DB (28), and NAStructuralDB (29). The latter two focus specifically on antibody/nanobody–antigen complexes, and thus do not contain *apo* antibody structures. All three databases are only available as downloads and are not maintained weekly, nor do they allow for direct searches, filtering, and analysis of antibody structures.

### Versioned train/test splits of ML-grade data enable fair and realistic comparison of ML models

In addition to collecting and annotating all publicly available antibody structures, we curate and clean a high-quality subset of SAbDab2 to create an ML-ready dataset. At launch, it includes 15 641 variable-region structures (1 739 *apo*; 13 902 *holo*, 1 245 of which involve multi-polymer antigens), corresponding to 5 301 unique SAbDab2 IDs (462 available in both *holo* and *apo* states, 4 158 *holo* only; 219 *apo* only; 388 with multi-polymer antigens).

To facilitate comparisons between ML models, we are also releasing standardised, versioned, backward-compatible train/test splits of this dataset, designed to mitigate against the data leakage concerns affecting the date-based splits currently prevalent in the literature

## Conclusion

SAbDab2 collects, curates, and annotates all antibody structures deposited in the PDB and presents them in a format readily accessible for computational analysis and ML applications. Building on the original SAbDab, which has served as a central resource for over a decade, SAbDab2 introduces improved support for rare and engineered antibody formats and constructs, and provides improved antigen annotations. Grouping identical numbered antibody sequences under a single SAbDab2 ID further enables systematic analysis of structural conformations, flexibility, and epitope diversity across structures.

In addition, SAbDab2 provides standardised, versioned dataset splits for antibody structure prediction and anti-body–antigen complex modelling. These datasets allow the community to consistently benchmark existing and emerging computational models.

SAbDab2 is updated weekly and can be accessed freely online under a CC-BY 4.0 license at https://sabdab2.opig.stats.ox.ac.uk. Standardised data splits of ML-grade data are available at https://zenodo.org/records/20083995, and updated versions will be made available once every six months.

## Methods

PDB entries were processed to identify numerable antibody segments using ANARCII (25). These segments were paired and classified, and antigens were assigned based on spatial proximity thresholds.

### Scraping the Protein Data Bank

Sequences (“pdb_seqres”), entries (“entries.idx”), and entry types (“pdb_entry_type”) were downloaded from the PDB file archive (https://files.wwpdb.org/). For all PDB entries classified as proteins we downloaded the PDBx/mmCIF files. Author-defined chain identifiers comprising numeric or alphanumeric characters were reassigned to avoid conflicts with SAbDab2’s multi-domain segment naming convention, in which chains containing multiple numerable domains are split.

### Identifying numerable segments with ANARCII

AN-ARCII (25), an antigen receptor numbering tool, was used to classify and number antibody sequences, including conventional antibodies, multi-domain antibodies, and Variable New Antigen Receptors (VNARs). The protein sequences were used as input for ANARCII. Sequences were initially evaluated in “unknown” mode, and those with an antibody score greater than 22 could be confidently classified as numerable antibody domains. Chains longer than 225 residues are reprocessed in “SCFV” mode to identify potential multi-domain architectures. Sequences classified as antibody chains by ANARCII but annotated as T-cell receptors (TCRs) by PDB entity titles were flagged for manual review. VNARs are identified based on an ANARCII score in “VNAR” mode exceeding both the “unknown” mode score and a threshold of 15, in combination with supporting metadata, and are always manually validated. Sequences with intermediate scores (15–22) in ANARCIIs “unknown” mode and containing conserved IMGT residues are also manually assessed. These conserved positions include cysteines at IMGT positions 23 and 104, a tryptophan at position 41, and a cysteine, phenylalanine, tryptophan, or tyrosine at position 118.

### Splitting numerable segments and antigens within author-designated chains

The identified numerable segments were aligned to the corresponding structurally resolved protein sequence. Gemmi (34) was used for processing all structures, and alignments are performed using the “pairwise2” function of BioPython (35). Unresolved and non-standard residues were stored.

Author-defined residue indices were overwritten with the IMGT numbering of the sequence. When an author-defined chains contained multiple domains, as identified by ANARCII, the chain was systematically split and renamed to reflect individual domains. For example, chains originally labeled “A” containing both heavy and light domains were reassigned as “A1” and “A2” based on their sequential order. PDBx/mmCIF data blocks were updated accordingly.

Author-defined chain identifiers were also reassigned when multiple entities— as defined in the PDBx/mmCIF file—were labelled with the same author chain ID. All entities were classified into the following categories: protein, peptide (protein chains shorter than 50 amino acids), peptide/nucleic acid, sugar, DNA, RNA, nucleotide, hapten, ion, chemical element, water, or other.

### Heavy and light chain pairing

Heavy and light domains within a PDB entry were paired when the distance between the *C*_*α*_ atoms at conserved IMGT position 104 is less than 22 Å. Structures in which this distance fell between 22 and 28 Å were manually evaluated. Pairing was not permitted when the heavy and light domains belong to different author-defined biological assemblies, unless overridden by maintainers.

### Classifying antibody type

Antibody instances were classified as “FV” (paired heavy and light chain), “FAB” (paired heavy and light chain with constant domains), “FAB+FC” (Fab with a Fc region), “SD-H” (single-domain heavy chain), “SD-L” (single-domain light chain), or “VNAR” (single-domain new variable antibody receptor). In addition, we annotated the higher-order construct in which these domains occur. An antibody instance can be part of engineered constructs such as single-chain Fvs (“SCFV”), diabodies (“DIABODY”), Dual Variable Domain (“DVD”) antibodies, Cross-Over Dual Variable (“CODV”) antibodies, single-chain diabodies (“SCDB”), or other antibody constructs (“OTHER”). Instances of the “FAB+FC” type can also be part of a full antibody construct (“FULLAB”; e.g. a traditional IgG). An scFv consists of one chain containing a single paired heavy and light chain. A diabody consists of two chains, each containing linked heavy and light domains, with pairing occurring between domains on opposite chains. DVD antibodies comprise two chains containing two variable domains of similar types (either heavy or light), with canonical pairing between equivalent domain positions across chains. CODV antibodies are simmilar to DVD antibodies, but employ cross-over pairing between non-equivalent domains on opposing chains. SCDBs consist of a single chain containing two heavy and two light domains, forming two paired FVs.

To assign “FAB” and “FAB+FC” classifications, we assessed the presence of constant domains in the resolved protein structure. Sequences containing numerable antibody segments were aligned against germline constant domain sequences from the IMGT (36) to identify these constant regions.

### Antibody–antigen pairing

Antigens were assigned to antibody instances based on spatial proximity to CDR residues. Only entities within the same author-defined biological assembly (when provided) were considered. Proteins and peptides were classified as antigens if any of their Cα or Cβ atoms lay within 7.5 Å of the *C*_*α*_ or *C*_*β*_ atoms of antibody CDR residues, as defined by the IMGT scheme. All other entity types (excluding water) were assigned as antigens if any atom lay within 7.0 Å of the same atoms on the antibody CDR residues. SAbDab2 allows antibodies to be antigens and multiple antibodies can bind the same antigen.

### Query the IMGT for gene annotations

Metadata stored in the PDBx/mmCIF files was systematically parsed. The IMGT knowledge graph (IMGT-KG) (37) was used to annotate heavy and light chains with gene information, where available.

### Defining SAbDab2 IDs

Processed antibody instances were assigned SAbDab2 IDs. A SAbDab2 ID uniquely identifies single- and paired-chain variable regions by their IMGT numbering. Consequently, a single PDB ID can contain multiple SAbDab2 IDs, and a given SAbDab2 ID can include antibody instances derived from several PDB entries. Individual entries may also encompass antibody instances of different formats or constructs and with distinct antigen assignments. SAbDab2 IDs consist of 10 characters and the prefix `sabdab2_(e.g. sabdab2_H0142L00YY)`. The first 5 characters identify the heavy chain and start with “H”. The last 5 characters identify the light chain and start with “L”. Null-identifiers (“H0000” and “L0000”) indicate the absence of a chain, and thus the SAbDab2 ID concerns a single-domain antibody.

### ML-grade data preparation and train/test splits

To construct the ML-grade data, we extracted individual antibody instances and any attendant antigen chains from X-ray and cryo-EM structures of non-pathogenic antibodies in SAbDab2 with ≤ 3.5 Å resolution. Antibody chains were cropped to the variable region (IMGT positions 1–128) and discarded if residue anomalies (missing non-oxygen backbone coordinate; unknown, ambiguous, or non-canonical residue identity) occured at positions 5–123. Anomalies outside this core were removed by trimming. Sequence-terminal anomalies were also trimmed from polymeric antigen chains (protein, peptide, DNA, and RNA types), and trimmed antigen chains were retained when they constituted a single contiguous, non-anomalous segment that spanned at least five residues. Antibody-type antigens were excluded.

Train/test splits were constructed based on sequence similarity (fraction of identical residues in covered span of longer sequence after pairwise alignment). Two splits were constructed: an “ab-split”, based on antibody similarity alonge, and an “ab-ag-split”, that also accounts for similarity among polymer antigens. For both splits, antibody instances were first grouped into separate base clusters on the CDRH3, concatenated CDRH123, and concatenated CDRL123 sequences (85% sequence identity thresholds for the “ab-split”; 95%, 100%, and 100% respectively for the “ab-ag-split”). Instances were then grouped into antibody clusters using connected-component clustering if they shared a base cluster of any type. For the “ab-split”, the resulting antibody clusters (connected components) were then allocated to train and test to achieve an approximate 8:2 ratio of antibody instances, while maintaining backward compatibility with previously published split versions.

For the “ab-ag-split”, we additionally created base clusters for polymeric antigen chains (40% sequence identity threshold for proteins; 80% for all other types). Antibody instances were then clustered together if they were members of the same antibody cluster (see above), or if any of their polymeric antigen chains shared an antigen base cluster. These resulting connected components were then allocated to train and test as before.

### Database implementation

SAbDab2 is implemented as a SQL database using the MariaDB management system, with query and access logic handled by custom Python implementation. The object-relational mapping between Python and SQL is implemented using the SQLModel library and SQLAlchemy toolkit, while querying and accessing the data itself is handled using an API implemented using the FastAPI framework. The SAbDab2 website presentation and search interface are implemented using the React framework and interact with the SQL back-end via FastAPI routes, as well as embedding the Molstar viewer (38) plugin for molecular visualisation.

## AUTHOR CONTRIBUTIONS

CMD conceptualised the study. HLC and OV processed the data, performed the research, and analysed the data. BHW and CRT implemented the database. HLC, OV, BHW, and CRT designed and implemented the website. CMD supervised the project. The manuscript was written by HLC and OV and reviewed by BHW, CRT, and CMD.

## ACKNOWLEDGEMENTS

We thank Dr Lewis Chinery and Dr Fergus Boyles for their extensive work on updating SAbDab over the years.

## Footnotes

1 https://sabdab2.opig.stats.ox.ac.uk/statistics

2 https://www.rcsb.org/news/feature/67edb35096cbbd16fc52edaf

